# Strategies to modulate zebrafish collective dynamics with a closed-loop biomimetic robotic system

**DOI:** 10.1101/831784

**Authors:** Yohann Chemtob, Leo Cazenille, Frank Bonnet, Alexey Gribovskiy, Francesco Mondada, José Halloy

**Affiliations:** Univ Paris Diderot, Sorbonne Paris Cité, LIED, UMR 8236, 75013, Paris, France; Department of Information Sciences, Ochanomizu University, Tokyo, Japan; Biorobotics Laboratory, School of Engineering, Ecole Polytechnique Fédérale de Lausanne, ME B3 30, Station 9, 1015 Lausanne, Switzerland

**Keywords:** collective behaviour, mixed societies, biohybrid systems, zebrafish, biomimetic robotics, closed-loop

## Abstract

The objective of this study is to integrate biomimetic robots into small groups of zebrafish and to modulate their collective behaviours. A possible approach is to have the robots behave like sheepdogs. In this case, the robots would behave like a different species than the fish and would present different relevant behaviours. In this study, we explore different strategies that use biomimetic zebrafish behaviours. In past work, we have shown that robots biomimicking zebrafish can be socially integrated into zebrafish groups. We have also shown that a fish-like robot can modulate the rotation choice of zebrafish groups in a circular set-up. Here, we further study the modulation capabilities of such robots in a more complex set-up. To do this, we exploit zebrafish social behaviours we identified in previous studies. We first modulate collective departure by replicating the leadership mechanisms with the robot in a set-up composed of two rooms connected by a corridor. Then, we test different behavioural strategies to drive the fish groups towards a predefined target room. To drive the biohybrid groups towards a predefined choice, they have to adopt some specific fish-like behaviours. The first strategy is based on a single robot using the initiation behaviour. In this case, the robot keeps trying to initiate a group transition towards the target room. The second strategy is based on two robots, one initiating and one staying in the target room as a social attractant. The third strategy is based on a single robot behaving like a zebrafish but staying in the target room as a social attractant. The fourth strategy uses two robots behaving like zebrafish but staying in the target room. We conclude that robots can modulate zebrafish group behaviour by adopting strategies based on existing fish behaviours. Under these conditions, robots enable the testing of hypotheses about the behaviours of fish.

## I. Introduction

Current research in engineering and science is seeking to bridge the gap between living and artificial systems to design new technologies. Building on biomimetic approaches, a stream of this research trend aims to build biohybrid systems that harness the advantages of both living and artificial systems [1]. Some of these biohybrid systems comprise robots and animals producing collective behaviours [2]. Here, we focus on experiments with autonomous robots that interact socially in closed loops with animals, that is, robots perceiving and reacting to animals and vice versa. A limited number of research studies deal with the difficult task of producing collective, close-loop animal-robot behaviours, especially controlling entire biohybrid systems to perform predefined tasks.

For example, Vaughan *et al.* [3] showed that a robot can gather ducks by acting like a sheepdog. Halloy *et al.* [4] demonstrated that multiple robots socially integrated and interacting in a closed loop with cockroaches could be used to control their decision-making mechanisms to reach predefined targets. Swain *et al.* [5] explored the interactions between a mobile robot and a shoal of Golden shiners by presenting the robot as a predator. Kawabata *et al.* [6] showed that a robot can modulate the behavioural reactions of a cricket. Kopman *et al.* [7] showed that fish adapt their behaviour to the tail-beating of a robotic lure and also that this response can be improved when the robotic tail beating is controlled, in closed loop, according to the movement of the fish. Landgraf *et al.* [8] showed that a mobile robot can interact with a shoal of Guppies and make recruitment. Shi *et al.* [9] showed that a robotic rat can modulate the behaviour of several laboratory rats with or without direct interactions. Donati *et al.* [10] showed that a dummy attached to a robotic arm can socially interact with the electric fish *Mormyrus rume*. Landgraf *et al.* [11] showed that a mobile robot with realistic eye dummies and natural motion patterns significantly improved its acceptance level among a shoal of Guppies. Stefanec *et al.* [12] showed that static robots simulating the presence of individuals by controlling their temperatures could interact and attract young bees. Worm *et al.* [13] showed that a robot moving randomly can interact with the electric fish *Mormyrus rume* by displaying prerecorded electric organ discharges (EODs) in answer to the EODs made by the live fish.

Cazenille *et al.* [14], [15] showed that a mobile robot capable of biomimetic behaviours can successfully integrate a group of zebrafish and produce collective behaviours. Bonnet *et al.* [16] showed that such mobile robots can modulate the collective decision-making of a group of zebrafish in a circular set-up. Papaspyros *et al.* [17] improved the integration of the robot under similar conditions by using a biomimetic model to drive it. Quinn *et al.* [18] showed that rats can release robot-rats from restrainers as they do for other conspecifics and can discriminate between different behaviours the robots expressed. Kim *et al.* [19] used a robotic arm with a zebrafish lure moving in three dimensions in an isolated tank connected to the fish tank only visually to show that, if the system is in a closed loop, some transfer learning can be observed. Bonnet *et al.* [20] showed that robotic fish and robotic bees integrated with their respective conspecifics could produce cooperation between the fish and the bees, respectively, in real time and over long distances.

While these previous studies succeeded in modulating animal behaviour with robots in closed loops, few used mobile robots on social animal groups, and even fewer used them with biomimetic trajectories. In studies involving fish, only Cazenille *et al.* [14], [15] used a biomimetic model to drive a fish robot. However, this work did not modulate fish behaviour, as Bonnet *et al.* [16] and Papaspyros *et al.* [17] achieved in a simpler environment.

Here, we study the modulation capabilities of fish robots integrated into zebrafish (*Danio rerio*) groups in a complex environment. We aim to achieve this by reproducing specific behaviours of collective movement, such as the leadership described by Collignon *et al.* [21] and collective spatial distribution [22]. This allows us to validate hypotheses made about the collective behaviour of zebrafish and gain a better understanding of zebrafish group decision-making. Leadership is an essential element in collective decision-making, as it reflects the will of one or a minority of individuals to initiate the movement of the group in a new direction. We have implemented theoretical leadership mechanisms in the robot to make it initiate collective departure in groups of fish.

We test different biomimetic behavioural strategies to drive the fish groups towards a predefined *target room* in a set-up composed of two rooms connected by a corridor. We test the hypothesis that, to drive the biohybrid groups towards a predefined choice, the robots have to adopt some specific, fish-like behaviours. Here, we test both a ‘bold’ behaviour with a robot performing collective departures towards the target and a ‘shy’ behaviour with a robot delaying collective departures from the target.

## II. Materials and Methods

### A. Ethics statement

All fish experiments were performed in accordance with the recommendations and guidelines of the Buffon Ethical Committee (registered to the French National Ethical Committee for Animal Experiments #40) after submission to the state ethical board for animal experiments.

### B. Animals and housing

We used groups of four or five adult, wild-type AB zebrafish (*Danio rerio*) in our experiments. The fish were 6–12 months old at the time of the experiments. We kept the fish under laboratory conditions: 27 *°*C, 500 *µ*S salinity with a 10:14 day:night cycle. The fish were reared in ZebTEC housing facilities and fed twice a day (Special Diets Services SDS-400 Scientific Fish Food). The water pH level was maintained at 7, and nitrites (NO^−2^) were below 0.3 mg l.

### C. Experimental setup

We used the set-up described in [15], [23], [22], [21] consisting of two rooms linked by a corridor (1 000× 1 000 × 100 mm^3^) made of white plexiglass (figure 1B) placed in a 1 200 × 1 200 × 300 mm^3^ experimental tank. The tank was filled with water up to a level of 60 mm. The whole set-up was exposed to diffused light and confined behind white sheets to isolate experiments and homogenise luminosity. We used the Fish-CASUs (Fish-Control-Actuator-Sensor-Unit) [24], [25], [23] to interact with the fish (figure 1A). They were composed of soft fish lures linked by magnetic coupling to mobile robots (FishBots) moving below the experimental tank. The fish lures had lengths of 4.5 cm that mimicked the morphology of the zebrafish by passively beating their tails when moving underwater. The floor of the aquarium was covered with a sheet of teflon to provide a smooth surface for the motion of the fish-lure. The FishBot robots were powered by two conductive plates, one glued to the bottom of the aquarium and one to the support on which the robot moved (figure 1C). An overhead acA2040-25gm monochrome GigE CCD camera (Basler AG, Germany) with a maximum resolution of 2 048 × 2 048 px and equipped with low distortion lenses CF12.5HA-1 (Fujinon, Tokyo, Japan) grabbed frames that were then processed to find the fish positions. A complementary fisheye camera (15 FPS, 640 × 480 px) was placed under the fish tank and was used to track the position of the robot. The tracking of the agents and the control of the robot were done using the CATS (Control and Tracking Software) framework [23].

**Fig. 1:**
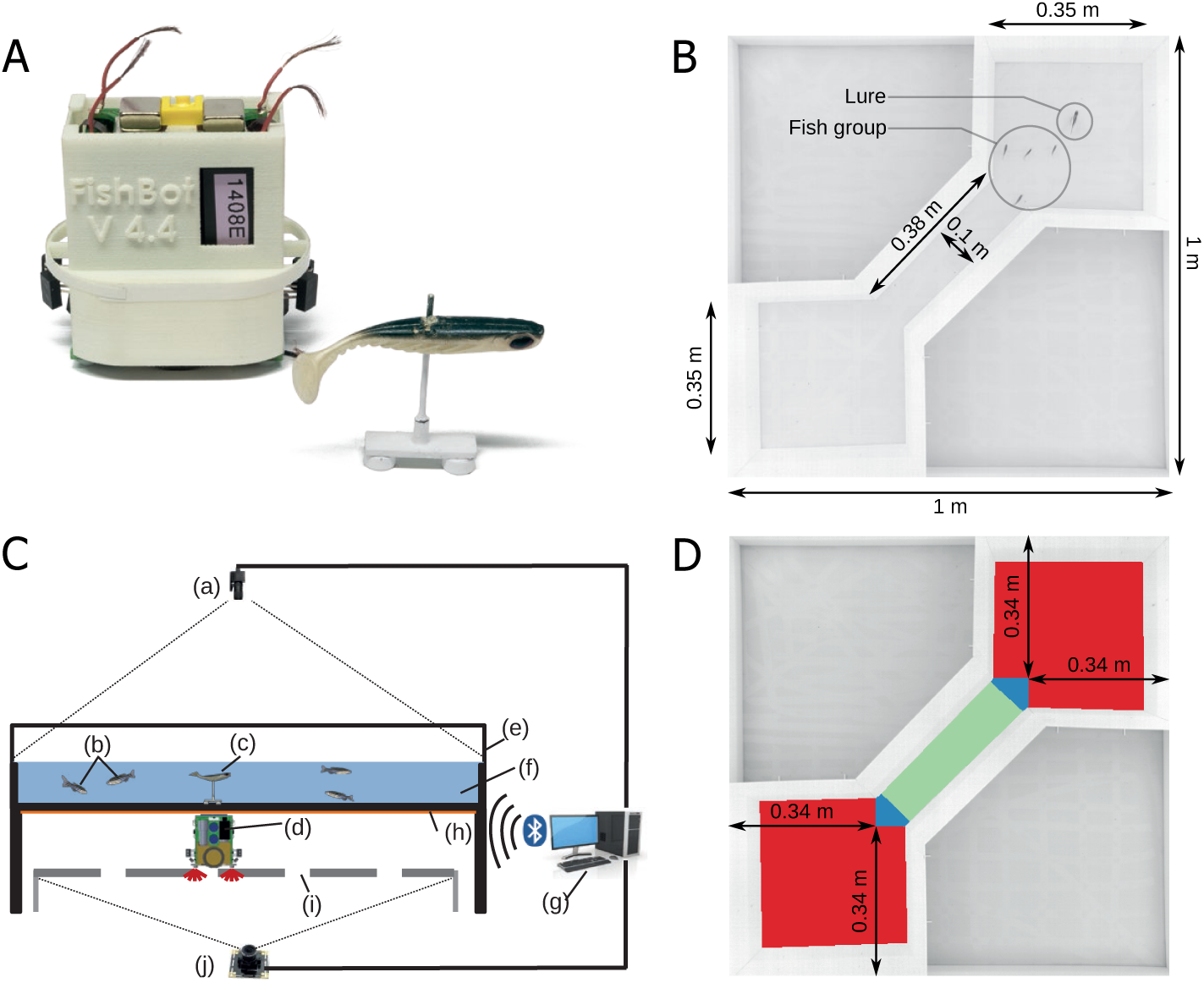
Experimental set-up. **A**: The FishBot [26], [24], [25] and a biomimetic lure used during the experiments. **B**: Experimental arena composed of two square rooms (350 mm × 350 mm at floor level) connected by a corridor (380 mm × 100 mm at floor level). **C**: Experimental set-up used during the experiments [26], [24], [25]. (a) Overhead camera used to track zebrafish. (b) Zebrafish. (c) Fish-lure. (d) FishBot moving under the aquarium and linked to the lure through magnetic coupling. (e) Fish tank of 1 000 × 1 000 × 250 mm. (f) Water layer 60 mm deep. (g) Computer processing camera frames and remote-controlling the robots via Bluetooth. (h) Conductive plate powering the mobile robot. (i) Perforated steel plate for powering the FishBot and to observe the FishBot LEDs from below. (j) 180 degree Fisheye camera to track the FishBot from below. **D**: Coloured zones of the arena corresponding to the three different types of behaviour of the robot [14]. When the robot is in one of the rooms (in red) or near the entrance of the corridor (in blue), it is driven by the biomimetic model presented in Sec. II-E. When the robot is in the corridor (in green), it drives straight through the corridor at a constant speed.

Each experiment was repeated 10 times and lasted 15 minutes. At the start of every trial, we let the fish occupy the set-up for 5 minutes with the robot not moving for acclimatisation.

### D. Data analysis

In addition to the online tracker included in the CATS framework [23], we acquired the identity of each agent using idTracker software [27] after the experiments. We obtained the positions *P* (*x, y, t*) of all agents at each time step Δ *t* = 1*/*15*s* for all experiments and built the trajectories of each agent. For each video, we manually selected which trajectory corresponded to the robot. Using the methodology developed in [21], we identified and detailed all collective departure. Using the agent positions, we also quantified the presence of fish in each part of the set-up.

### E. Perception-based behavioural model

To control the FishBots, we used the multilevel behavioural model presented by Cazenille *et al.* [14], which is derived from the model developed by Collignon *et al.* [28] and can simulate the collective behaviour of zebrafish in a finite environment. This model relies on two levels of control: a low level dealing with the movement patterns and a high level building the trajectories. Both the levels are biomimetic and context-dependent according to the different behaviour expressed by the fish and according to the geometry. In large environments, that is, the rooms in this set-up, the robot followed a perception model, while in narrow spaces, that is, the corridor, it followed a straight line at a constant speed.

This perception model (see [14]-III-A) is inspired by the probabilistic model described in [28]. The agents update their position vectors *X*_*i*_ with a velocity vector *V*_*i*_:

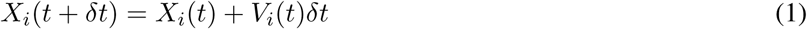

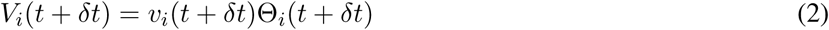

where *v*_*i*_ is the linear speed of the *i*^*th*^ agent and Θ_*i*_ is its orientation. The linear speed *v*_*i*_ of the agent is randomly drawn from the experimentally measured instantaneous speed distribution.

The Θ_*i*_ orientation is sampled from a probability density function (PDF) calculated as a mixture distribution of von Mises distributions centred on the stimuli perceived by the focal agent. The influence of other agents is taken into account.

We numerically calculated the cumulative distribution function (CDF) matching this final PDF by performing a cumulative trapezoidal numerical integration of the PDF in the interval [−*π, π*]. The model draws a random direction Θ_*i*_ in this distribution by inverse transform sampling. The agent’s position is updated according to this direction and linear speed.

## III. Defining different robot behaviour modulation strategies

Figure 2 shows the description of the algorithm run by the robot for the initiation experiment E2. The goal was to influence the group of fish to move from one room to another. The robot initiated collective departure from the *starting room* towards the *target room*.

**Fig. 2:**
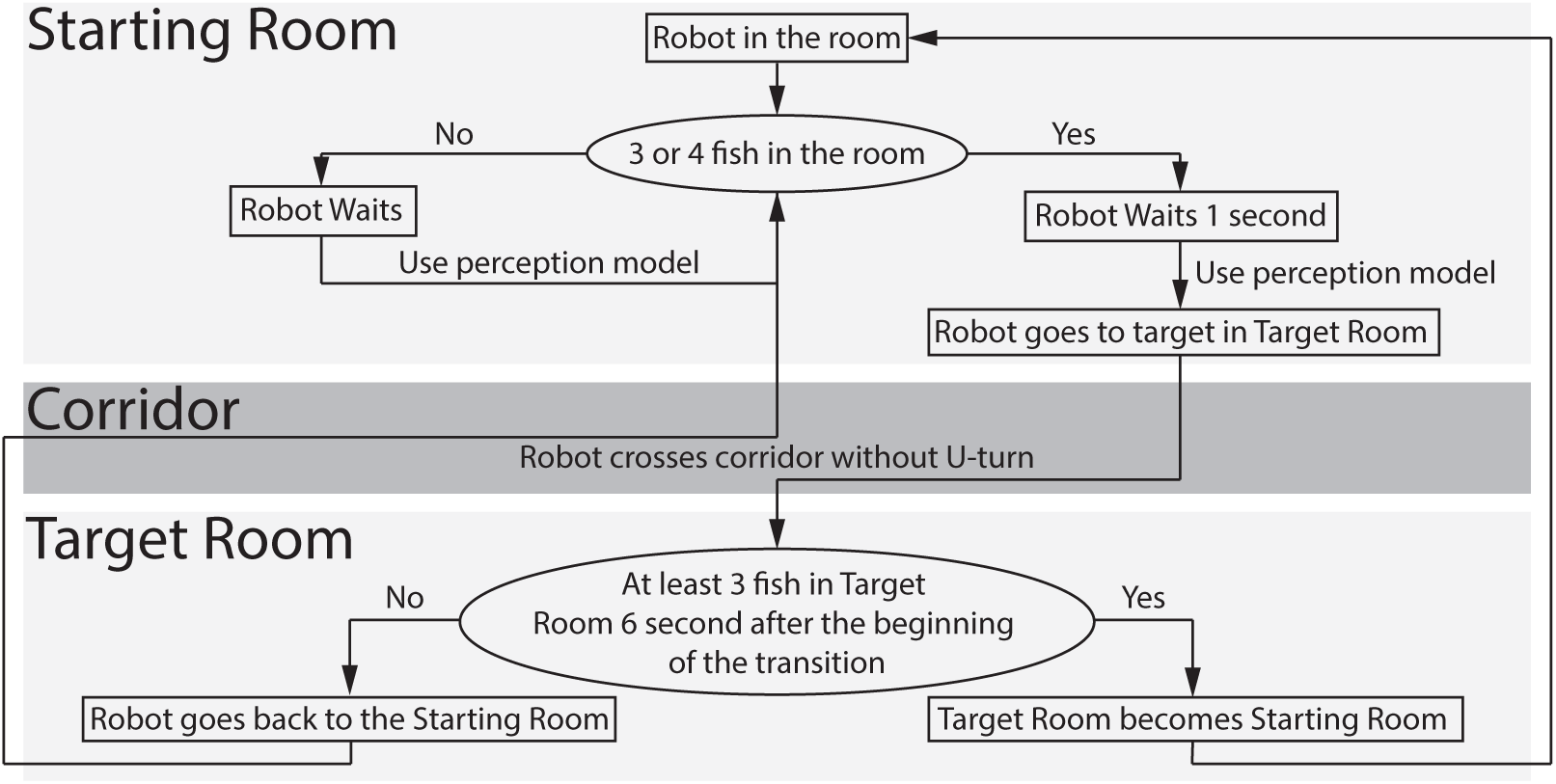
Initiation behavioural strategy of the robot. The behavioural model of the robot is a mix of biomimetic initiation behaviour and artificial behaviours developed to enhance the effect of the initiation strategy.

The robot had two phases in the *starting room*: when it waited for the group of fish and when it started a transition. When it waited, the robot stayed in the *starting room* using the biomimetic perception model described in Sec. II-E. If three fish or more were detected in the *starting room*, the robot turned in the room over the course of one second, using the model to attract the fish.

After that, the robot reached a target put in the *target room*. Six seconds after the beginning of the transition, the controller checked the number of fish in the *target room*. The six seconds delay corresponded to the maximum delay observed for the fish to make a transition. If at least three fish were detected in the room, the *target room* became the *starting room*, and the algorithm started again. If not, the robot went back to the *starting room*.

Ten trials were done with the robot following this behaviour over 15 minutes with four fish.

Experiment E1 was the biological reference case, involving 10 groups of five wild-type zebrafish and no robot. In experiment E3, we used the initiation behaviour described above to modulate the presence of four fish between the two rooms with one robot. We wanted to attract the fish to one room more than the other, so the *target room* and the *starting room* defined in the initiation behaviour were set at the beginning of each of the 10 trials and did not change during the trial. For experiments E4 and E5, either one or both robots used the following attraction behaviour on a group of four fish: the robot was driven by the biomimetic perception model but could only navigate inside the *target room*. The *target room* changed between each of the 10 trials and was selected randomly. This corresponded to the behaviour of a fish with a strong preference for one room and no exploration behaviour. Experiment E6 was a mix between experiments E3 and E4 with two robots and four fish. One robot used the initiation behaviour and the other the attraction behaviour. The *target room* was the same for the two robots and was selected randomly at the beginning of the 10 trials. All the trials lasted 15 minutes.

**TABLE I:** Strategies of modulation. The **E1** experiment is the biological reference case, 10 trials of five wild-type zebrafish groups and no robot. The **E2** experiment involves 10 trials of four wild-type zebrafish and one robot driven by the initiation strategy. The **E3** experiment involves 10 trials of four wild-type zebrafish and one robot driven by the initiation strategy. The **E4** experiment involves 10 trials with a robot driven by attraction behaviour and 4 zebrafish. The **E5** experiment is the same as **E4** but with two robots with fish-like behaviour. The **E6** experiment mixes these two strategies and uses two robots, one initiating and the other attracting.

| Label | Experiment | Number of agents | Robot strategy |
| --- | --- | --- | --- |
| E1 | Reference AB zebrafish only | 5 fish + 0 robot | – |
| E2 | Initiation of transitions | 4 fish + 1 robot | Biomimetic initiation without target room |
| E3 | Modulation by initiation | 4 fish + 1 robot | Biomimetic initiation to target room |
| E4 | Modulation by attraction | 4 fish + 1 robot | Fish behaviour & stay in target room |
| E5 | Modulation by attraction | 4 fish + 2 robots (R1,R2) | R1 & R2 stay in target room |
| E6 | Mixed initiation and attraction | 4 fish + 2 robots (R1,R2) | R1 biomimetic initiation & R2 stays in target room |

## IV. Results

### A. Initiation of collective departure

In a previous study [21], we showed that zebrafish present a shared leadership: each fish has the same ability to initiate a collective movement, which our experiments demonstrated with the action of initiating a transition from one room to the other. However, zebrafish groups exhibit heterogeneity, ranging from groups in which leadership-sharing is egalitarian to groups in which a single fish monopolises the leadership. This difference is explained by the greater mobility of these individuals. Here, we emulated this behaviour by forcing the robot to perform as many transitions as possible.

We validated with experiment E2 that the robot can, in fact, lead the group with this behaviour. Here, every time the robot was with the most fish in one chamber, it exited the room to go to the *other room* (Sec III).

Our results showed (figure 3A) that, in many trials with different groups of fish, the robot initiated a transition of the group of fish more often than all the other fish in the group (7 / 10) or all the fish except one (3 / 10).

**Fig. 3:**
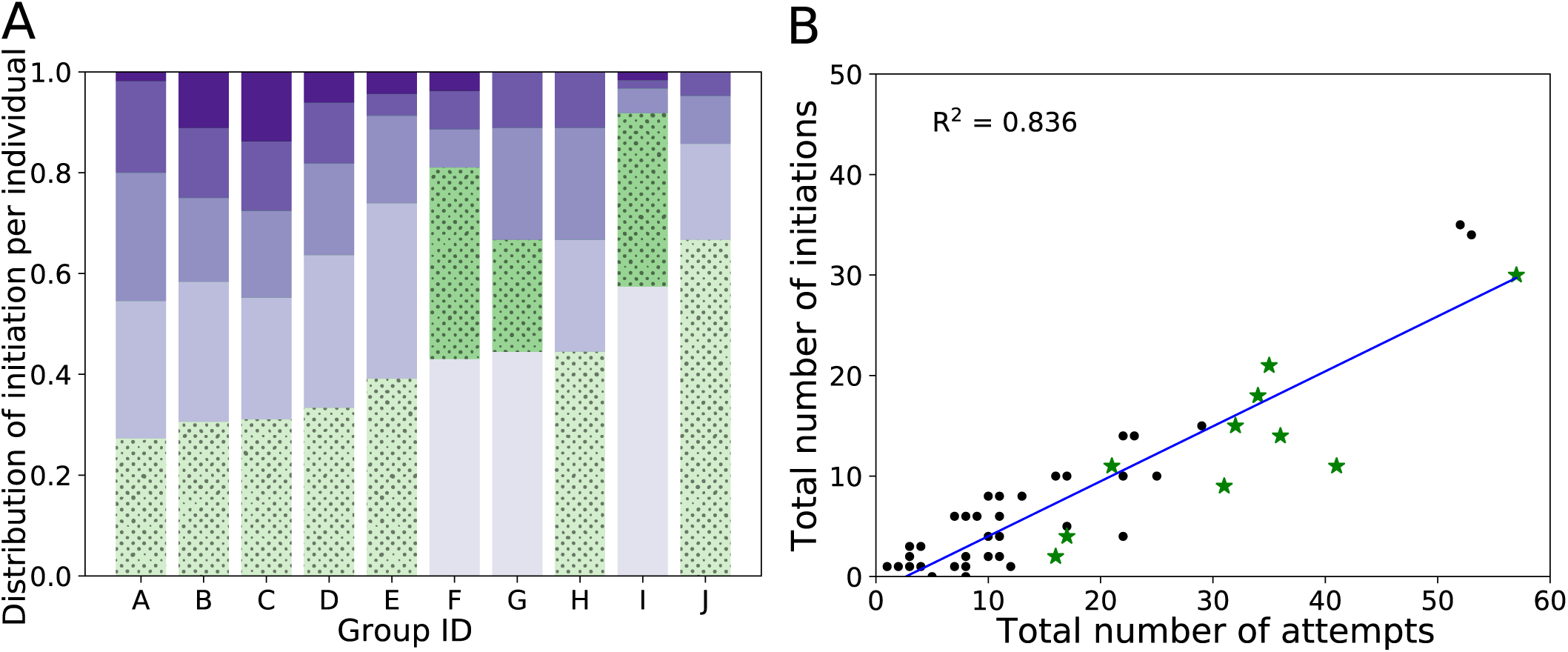
Robot initiation of transition towards the *target room*. (A) Frequency of initiation according to the intra-group ranking of the fish for initiation. For each group, we calculate the proportion of group departure that each individual initiated, as in [21]. The fish are then ranked according to this frequency. The robot is indicated in dotted green. (B) The total number of collective departures initiated in relation to the total number of attempts made by fish, in black, and by the robot, in green. The data show that the number of initiations is directly proportional to the number of attempts. Each fish has the same probability of success to initiate a collective departure when attempting. The robot, even with more initiations than most fish, has a similar probability of success.

As in [21], a linear regression shows that the total number of initiations of each fish is linearly correlated to the total number of attempts performed (figure 3B). This means that the success of fish that initiate more group starts than their congeners can be explained simply by the fact that they tried to exit the room more often [21], not by their physical traits, such as size or sex. In our experiment, we applied this hypothesis to the robot behaviour, and figure 3B shows that the ratio of attempts and successes of the robot initiating collective departure, represented by green stars, corresponds to the fish. We also compared the distribution of the number of initiations by individuals between the reference experiment E1 and experiment E2 using a t-test and did not observe any significant difference (p-value = 0.3880). The number of initiations performed by the robot was therefore not excessive compared to that observed in fish. This experiment demonstrated that we could reproduce zebrafish initiation behaviour as described in [21] with our robot during long trials, even using the robot as a strong leader. By comparing the robot trajectories in figure 4 during the initiation phases with a fish under similar conditions, we observed that the robot followed a trajectory with linear speeds comparable to those observed in fish. In the example figure 4B, we observed that fish used the same trajectories as the robot with a one seconds delay. Also, the fish exited the rooms in groups.

**Fig. 4:**
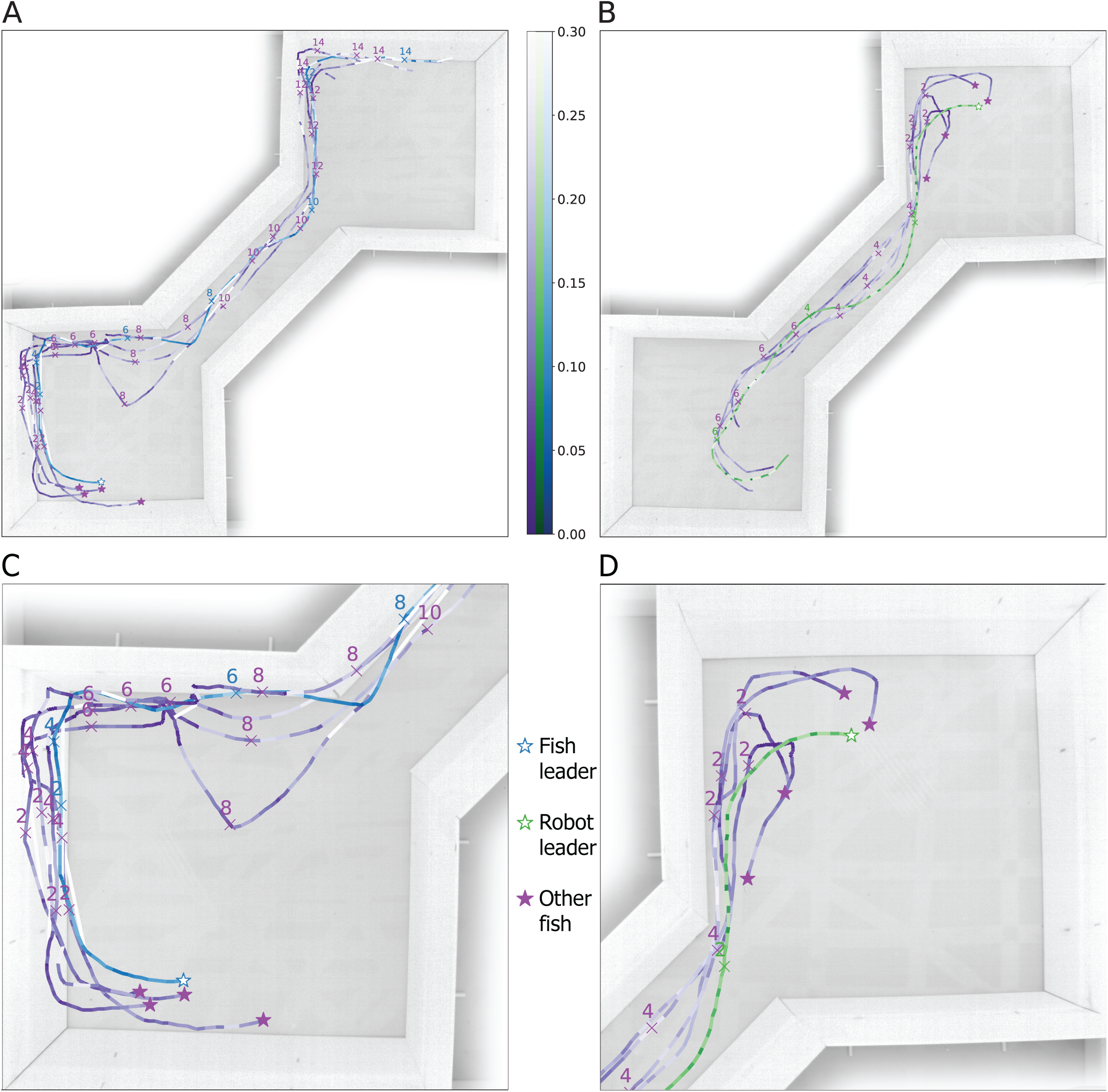
Examples of individual trajectories during a collective departure event (A) with a fish as a leader from the fish control experiment E1 and (B) with the robot as the leader, as in experiment E2. (C) and (D) are zooms of (A) and (B), respectively. The trajectory of the leader is in green for the robot, in blue for the fish and in purple for the other fish. The colour scale indicates the linear speed of the agent in *m/s*. A star indicates the beginning of the trajectory, with a white centre for the leaders. Crosses on the trajectory indicate the time step in seconds. By comparing the trajectories of the robot and fish, we observe that the robot succeeds in reproducing the initiating behaviour.

### B. Modulation of the spatial distribution

We wanted to modulate the collective decisions of the fish with the robot and change the time spent by the fish in each room. We considered two main strategies: first, using the robot as a leader to create a collective decision of exiting one room to a *target room*, and second, attracting the fish in the *target room* using the robot as an active lure. We also combined these strategies in an experiment with two robots, one the leader and one the attractor.

Experiment E1 was our reference with five fish and no robot. We demonstrated that the fish did not have a preference for either room. We measured the time spent in each room (figure 5A). We looked for differences between the two distributions using a Wilcoxon test and observed no significant difference (p-value = 0.8886). Without robot interference, the dfish were in each room the same amounts of time. We also observed that they spent less time in the corridor, so this part of the set-up was considered a transition zone.

**Fig. 5:**
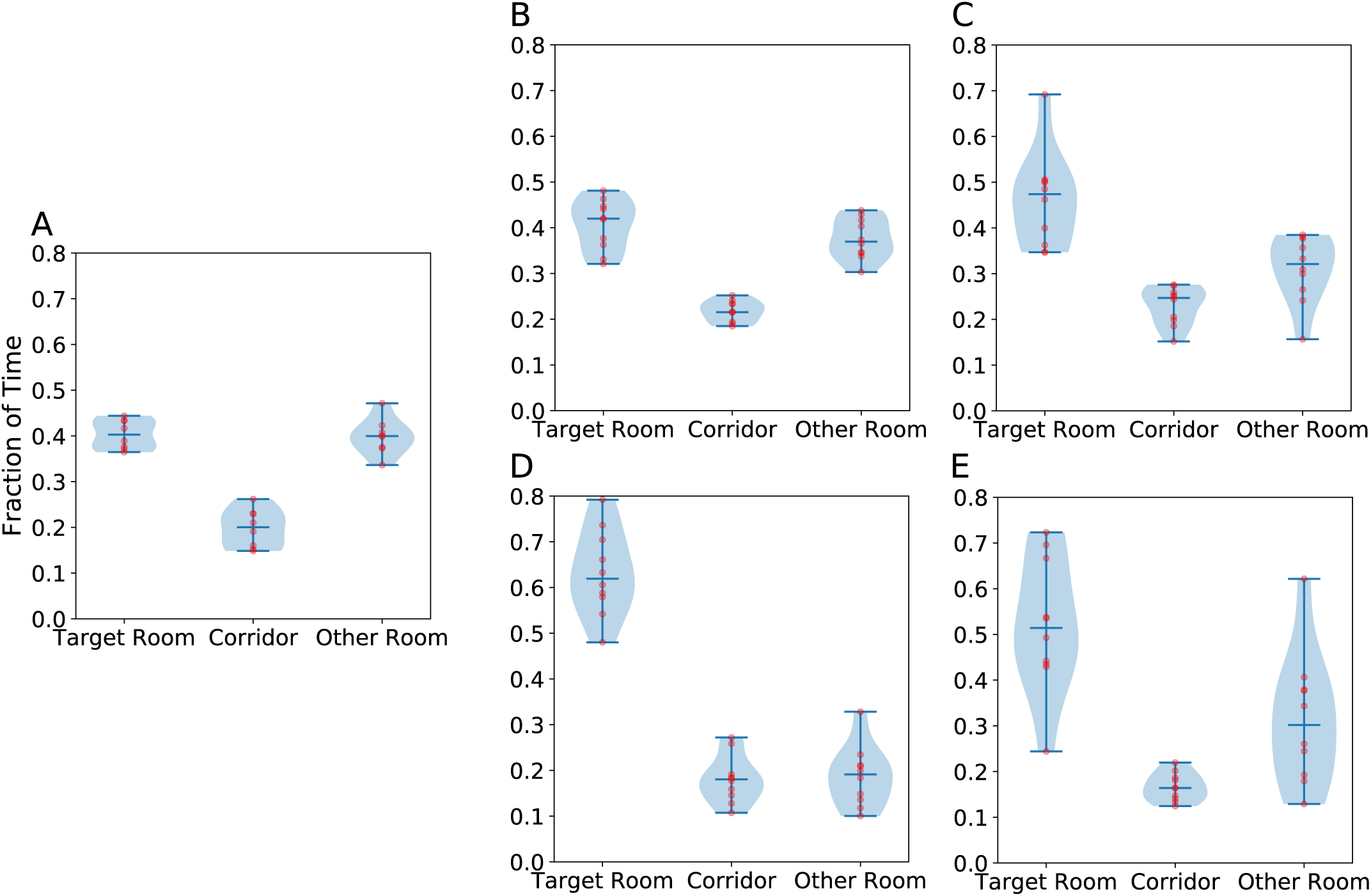
Fraction of time spent by all groups in the five experiments: **E1** Use of space for only five fish in the different parts of the two-room set-up (A), **E3** for groups comprising four fish and one robot acting as a leader and initiating transitions towards the *target room* (B), **E4** groups of four fish and one robot acting as an attracting lure in the *target room* (C), **E5** groups of four fish and two robots acting as attracting lures in the *target room* (D), **E6** groups of four fish and two robots, one acting as an initiating leader and the other an attracting lure (E). A Wilcoxon test shows no significant difference between fish presence in the rooms in (A), as fish spend the same amount of time in both rooms and transit rapidly through the corridor between the rooms (p-value = 0.8886) and (B) (p-value = 0.3329) when there is significant difference of the spatial distribution in (C) (p-value = 0.0284), (D) (p-value = 0.0051), (E) (p-value = 0.0367).

In experiment E3, we used the initiation behaviour tested in E2 to modulate the fish spatial distribution during a 30 minutes trial. Figure 5B corresponds to experiment E3, in which the robot acted as a leader and initiated transitions towards the *target room*. There was no significant difference between the time spent in each room (p-value = 0.3329). When using the initiation behaviour, the robot failed to modulate the spatial distribution. However, looking at the initiation successes in each of the rooms (figure 6), we can see that the robot succeeded in initiating collective departures to the *target room* (except for one group) (figure 6A) and did not lead in the other direction (figure 6B), as expected. The failure of the modulation of spatial distribution could therefore be explained by the fact that zebrafish have short residence times [22]. The collective departures of the robot may therefore have occurred in a timeframe close to that executed by the fish themselves.

**Fig. 6:**
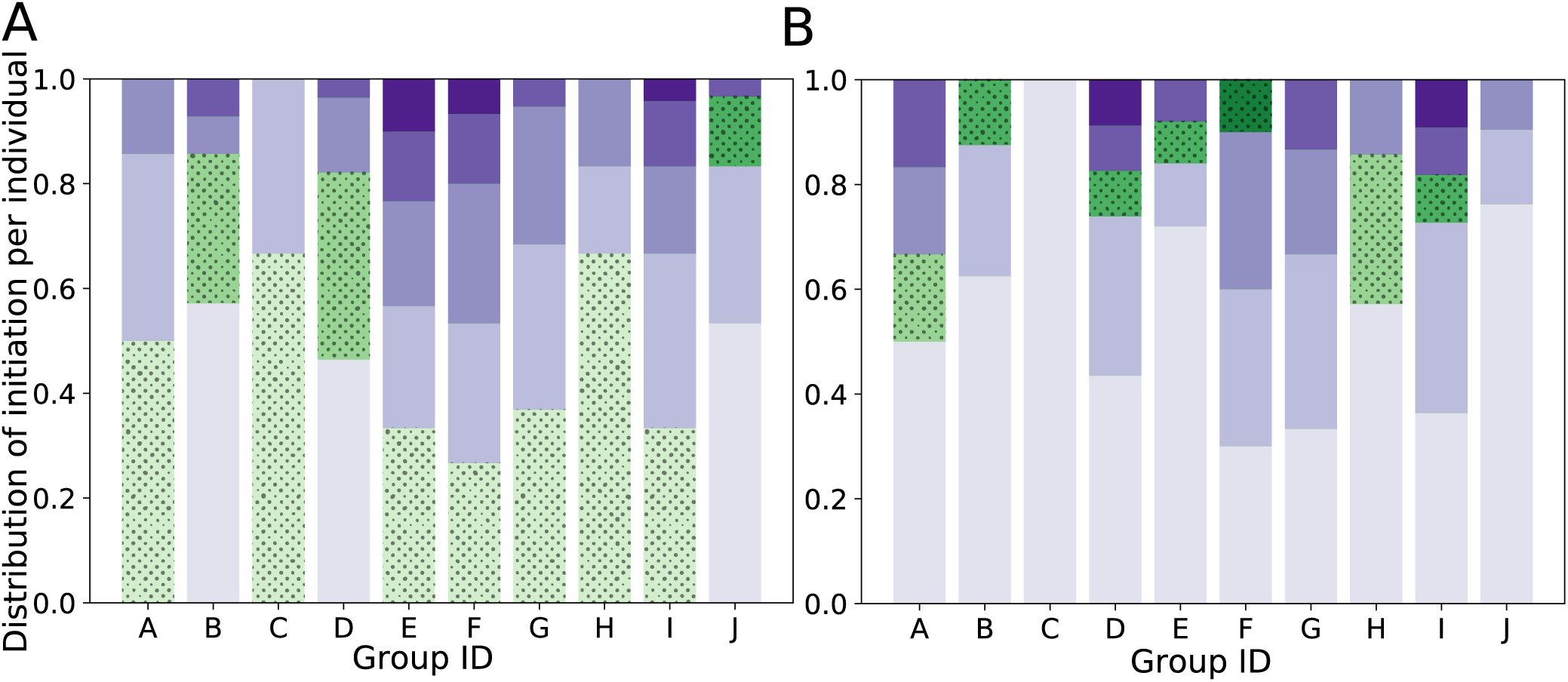
Proportion of departures initiated by each agent in experiment E3. The robot is indicated in dotted green. (A) To the target chamber. The robot is a leader. (B) From the target chamber. The robot is not the leader. The results indicate that the modulation of the transitions towards the *target room* works (A), although the residence time in the *target room* is not significantly changed (figure 5B).

Another strategy to modulate the spatial distribution is therefore to delay collective departures thanks to an agent that represses any exit from the *target room*.

In experiment E4 (figure 5C), the robot acted as a lure by swimming only in the *target room* during all the trials. We observed that fish stayed more in the *target room* (p-value = 0.0284) with a median of 0.45 against 0.31 in the *other room*. However, the variability of the time spent by the shoal in each room was quite high between trials, with values in the *target room* ranging from 0.3 to 0.7. This behaviour significantly modulated the fish, but some groups of fish seemed less attracted by the robot.

In experiment E5 (figure 5D), we used two robots with the same behaviour as in E4. The difference between the medians was higher than in E4 (0.63 against 0.18), and the variability was reduced between trials. The effect observed with one robot was amplified with two robots, and the influence of the robots along all groups was constant.

From the results above, we combined experiments E3 and E4. In experiment E6 (figure 5E), one robot acted as a leader to encourage the group to exit the *other room* while the other robot acted as a lure to delay the group from exiting the *target room*. There was a significant difference between the occupation of the two rooms (p-value = 0.0367). The medians were similar to the ones found in E4 (figure 5C), but the variability was even higher than in E4. Instead of being complementary, the initiation behaviour had a diminishing effect on the attraction behaviour.

## V. Discussion

Previously, we have analysed the collective behaviours of small zebrafish groups in structured environments composed of two rooms connected by a corridor [22], [21]. This set-up was designed to study how zebrafish groups navigate between the two rooms and how they make decisions to switch from one room to the other. In parallel, we developed an autonomous robot capable of zebrafish biomimetic behaviours that can be socially integrated into small zebrafish groups [29], [14]. We further showed that a small number of those robots can modulate the collective behaviour of mixed zebrafish and robot groups in a circular set-up [16].

Here, we further explored the possibilities of modulating the behaviour of small zebrafish groups by using the same, previously integrated robot. We tested several strategies inspired by the biological results obtained by Collignon *et al.* [21] and Seguret *et al.* [22].

We used the robot to modulate collective departure over a period of about 15 minutes. The objective was to test the hypothesis that an individual with bold behaviour could exert a form of leadership by initiating transitions between the two rooms more often than other members of the fish group. We used the same behaviours as [14], but we added a behavioural rule that made the robot initiate a room transition as often as possible. Under these conditions, the robot could induce transitions more often than most of the fish. By looking at the ratio between exit attempts and initiations of movement from one room to another, we observed that the robot followed the same linear relationship as that observed in fish. These two results are consistent with what has been observed in zebrafish groups alone [21]. It also confirms the hypothesis that ‘bold’ behaviour is sufficient to explain the variations observed in the distribution of leadership of *Danio rerio*. This conclusion goes further than our previous work with autonomous robots [14], [30], [15]. Indeed, the initiations of collective departures have never been explicitly simulated by a fish robot before. In this study, we reproduced this process in a way similar to what can be observed in fish [21].

We then used the robot to bias the residence time of fish between the rooms.

Once we showed that the robot could induce group transitions between the rooms, making the robot choose one of the two rooms as the *target room* was simple. The hypothesis was that, by making more transitions towards that *target room*, the fish robot could bias the fish group presence density in favour of the *target room*. The experiments showed that this did not work, as the presence of the fish groups remained equivalent in both rooms. This may be due to the fact that zebrafish are highly dynamic fish with short residence times that keep moving in their environments, never setting long in one room [22], [21]. As soon as they arrived in the *target room*, they did not stay long and made another transition towards the *other room*.

We tested another strategy to bias the residence time of the groups in favour of a *target room*. The fish robot made a choice and remained in the *target room* to become an attracting lure for the fish group. In this experiment, the results showed that, indeed, the fish groups spent more time in the *target room*. The luring strategy of using an individual (the fish robot) to attract the group is thus more efficient than inducing group transitions towards the *target room*. Moreover, this strategy was simpler in terms of behavioural program.

The recruitment effect can be enhanced by adding more robots behaving as robot-fish duets in the *target room*. In this case, the bias in favour of the target is maximal compared to the other strategies we tested in this study.

If we try to mix the two types of strategies, transition initiations and attraction, we see that the bias in favour of the target is low. In addition, the dispersion of the results showed that the group of fish was disturbed by this strategy. This disturbance was probably due to the mobility of the robots, which are not as fluid as that of fish.

This study dealt with groups comprising small numbers of zebrafish. We know that such small groups are more socially unstructured and less cohesive than larger groups. In this configuration, we chose the maximum difficulty to try to modulate the groups of fish with robots. Some of these strategies therefore allowed us to significantly bias the spatial distribution of fish groups over a long period of time by using robots moving with them. This was done previously in cockroaches [4] with robots selecting specific shelters. In fish, Bonnet *et al.* [16] succeeded in biasing the swimming direction of fish using the same robots. However, the environment used, a circular corridor, and the objective were both simpler than in our study. In the two-room set-up, the robots could not move according to a predefined path as in Bonnet *et al.* [16] but had to use a behavioural model to navigate the environment and interact with fish. This made it more complex to carry out modulation, but it allowed us to better understand decision-making within social fish groups.

Whether on collective starts or spatial distribution, we have shown that it is possible to modulate the behaviour of social fish with autonomous, biomimetic robots using strategies based on behaviours observed in animals. Our methodology allows us to test hypotheses about the functioning of specific collective behaviours. Other behaviours could therefore be tested with this same methodology.

## Acknowledgments

This work has been funded by the EU-ICT project ‘ASSISIbf’, no. 601074. The funders had no role in the study design, the data collection and analysis, the decision to publish, or the preparation of the manuscript. There was no additional external funding received for this study. The authors declare no conflicts of interest.

